# Human-lineage-specific genomic elements: relevance to neurodegenerative disease and *APOE* transcript usage

**DOI:** 10.1101/2020.04.17.046441

**Authors:** Zhongbo Chen, David Zhang, Regina H. Reynolds, Emil K. Gustavsson, Sonia García Ruiz, Karishma D’Sa, Aine Fairbrother-Browne, Jana Vandrovcova, International Parkinson’s Disease Genomics Consortium (IPDGC), John Hardy, Henry Houlden, Sarah A. Gagliano Taliun, Juan Botía, Mina Ryten

## Abstract

Knowledge of genomic features specific to the human lineage may provide insights into brain-related diseases. We leverage high-depth whole genome sequencing data to generate a combined annotation identifying regions simultaneously depleted for genetic variation (constrained regions) and poorly conserved across primates. We propose that these constrained, non-conserved regions (CNCRs) have been subject to human-specific purifying selection and are enriched for brain-specific elements. We find that CNCRs are depleted from protein-coding genes but enriched within lncRNAs. We demonstrate that per-SNP heritability of a range of brain-relevant phenotypes are enriched within CNCRs. We find that genes implicated in neurological diseases have high CNCR density, including *APOE*, highlighting an unannotated intron-3 retention event. Using human brain RNA-sequencing data, we show the intron-3-retaining transcript/s to be more abundant in Alzheimer’s disease with more severe tau and amyloid pathological burden. Thus, we demonstrate the importance of human-lineage-specific sequences in brain development and neurological disease. We release our annotation through vizER (https://snca.atica.um.es/browser/app/vizER).

## INTRODUCTION

Humans are perceived to be particularly vulnerable to neurodegenerative disorders relative to other primates on both a pathological and phenotypic level^1–5^. This is exemplified in Alzheimer’s disease, in which a similar phenotype is not seen in ageing non-human primates, nor are the characteristic neurofibrillary tangles on pathological examination^1,6^. Likewise, Parkinson’s disease does not naturally occur in non-human primates, whose motor deficits do not respond to levodopa administration and a Lewy body pathological burden is not present^5,7^. This has led to the hypothesis that the same evolutionary changes driving encephalisation which have steered the development of characteristic human features may predispose to disorders that affect the brain^2,5,6^. In the case of Alzheimer’s disease, it is postulated that the accelerated evolution of intelligence, brain size and aging predispose to selective advantages, which in later life, have deleterious effects on cognition through the very same pathways^8^. Therefore, identifying the genomic changes unique to the human lineage may not only provide insights into the evolution of human-lineage-specific phenotypic features, but also into the pathophysiology underlying uniquely human diseases.

Previous studies attempting to identify human-lineage-specific variation and functional elements in the human genome have focused on genomic conservation as calculated by aligning and comparing genomes across species. But, conservation measures alone do not fully identify regions with evidence of human-specific purifying selection. This is because a large part of the genome is evolving neutrally and sufficient phylogenetic distance is required to detect these changes^9^. Furthermore, alignment methods do not reliably detect substitutions that preserve function^9^. Conversely, some genes such as those implicated in immune system function may be subject to rapid evolutionary turnover even among closely-related species^9^. For these reasons, analysing conservation alone has limited capacity to capture human-specific genomic elements^9^.

The increasing availability of whole genome sequencing (WGS) has opened new opportunities to address this issue. Using intra-species whole-genome comparisons^10,11^, we are better able to appreciate sequence differences between individuals of the same species, and identify genomic regions in humans containing significantly fewer genetic variants than expected by chance, designated as constrained genomic regions. This form of analysis, which is based on the assumption that most selection is negative or purifying (i.e., those which remove new deleterious mutations), has been crucial for classification of exonic variation and attribution of pathogenicity^12^. However, many genomic regions would be expected to be both constrained and conserved; such regions have been maintained by natural selection across species, including humans. This means that metrics reflecting constraint alone cannot identify human-specific elements as the same regions could also be conserved in other species.

This has led previous analyses to combine these metrics of sequence constraint and conservation to identify genomic regions with evidence for human-specific selection^13,14^. Ward and Kellis successfully applied this approach to demonstrate that a range of transcribed and regulatory nonconserved elements showed evidence of lineage-specific purifying selection^14^. However, this analysis was limited by the availability of WGS data and metrics on human genetic variation were derived from the 1000 Genomes pilot data, which sequenced with only two to six times coverage^15^. Advances in sequencing technology have increased the feasibility of deep sequencing of human populations leading to a much more detailed understanding of genetic variation between humans^10^. In fact, the recent sequencing of the genomes of 10,545 human individuals at a coverage of 30 to 40 times identified 150 million single nucleotide variants of which 54.7% had not been reported in dbSNP^16^ or the most recent phase 3 of the 1000 Genomes Project^17^. The availability of this information has already enabled more accurate identification of relatively constrained regions of the genome, which has led to the development of the context dependent tolerance score (CDTS)^11^. CDTS is derived from estimating how the observed genetic variation compares to the propensity of a nucleotide to vary depending on its surrounding context using the high-resolution profiles determined from deep sequencing data^11^. Yet, this information has not been combined directly with improved conservation data to identify regions with evidence for human-specific selection.

In this study, we make full use of these resources to develop a novel, granular genomic annotation which efficiently captures information on intra-species constraint and inter-species conservation simultaneously and identifies constrained, non-conserved regions (CNCRs). We use this annotation to test the hypothesis that CNCRs are not only specific to the human lineage, but given the encephalisation of humans, that CNCRs will be enriched within brain-specific functional and regulatory elements as well as risk loci for neurological disease. We show that these regions are enriched for SNP-heritability for a range of neurological and psychiatric phenotypes. Furthermore, by calculating CNCR density within the boundaries of known genes, we develop a gene-based metric of human-specific constraint. This analysis highlights *APOE* and leads to the identification of an intron-3 retaining transcript of *APOE*, the usage of which is correlated with Alzheimer’s disease pathology and *APOE*-ε4 status. This approach provides direct support for the role of human-specific CNCRs in brain development and complex neurological phenotypes.

## MATERIALS & METHODS

### Generation of an annotation for the identification of CNCRs

We generated a combined annotation to capture information on intra-species constraint and interspecifies conservation simultaneously, using CDTS together with phastCons20 scores (**Figure 1**). The previously-validated map of sequence constraint (http://www.hli-opendata.com/noncoding) generated using 7,794 whole genome sequences^11^ was used to assign a single CDTS score to each non-overlapping 10 base pair (bp) region throughout the genome (build GRCh38, 248,925,226 bins). The phastCons20 score, which calculates the likelihood ratio of negative selection based on the total number of substitutions during evolution of an element between species^18^, was used as a measure of inter-species conservation (http://hgdownload.cse.ucsc.edu/goldenPath/hg38/phastCons20way/)^18^. PhastCons20 was used as it compares the human genome to the genomes of less divergent species (16 other primates and three mammals). For each 10bp bin labelled with a CDTS value, we assigned the corresponding mean phastCons20 score. Bins without a conservation score due to insufficient species in the alignment were not considered (0.218% of the genome). For the remaining 248,381,744 bins, we ranked both CDTS and mean phastCons20 scores across the whole genome such that the highest ranks represented the most constrained and conserved regions respectively. We calculated the log2 ratio of the rank of constraint to the rank of conservation for each 10bp bin (termed constrained, not conserved score, **CNC score**). This resulted in scores with a distribution centred at 0 signifying no fold change between the ranks of the two metrics (**Supplementary Figure 1**). Finally, we defined CNCRs as genomic regions that were among the 12.5% most constrained, with a CNC score of ≥ 1 (i.e. a two-fold higher ranking in constraint than conservation). We use this definition for CNCRs throughout this study.

**Figure 1.**
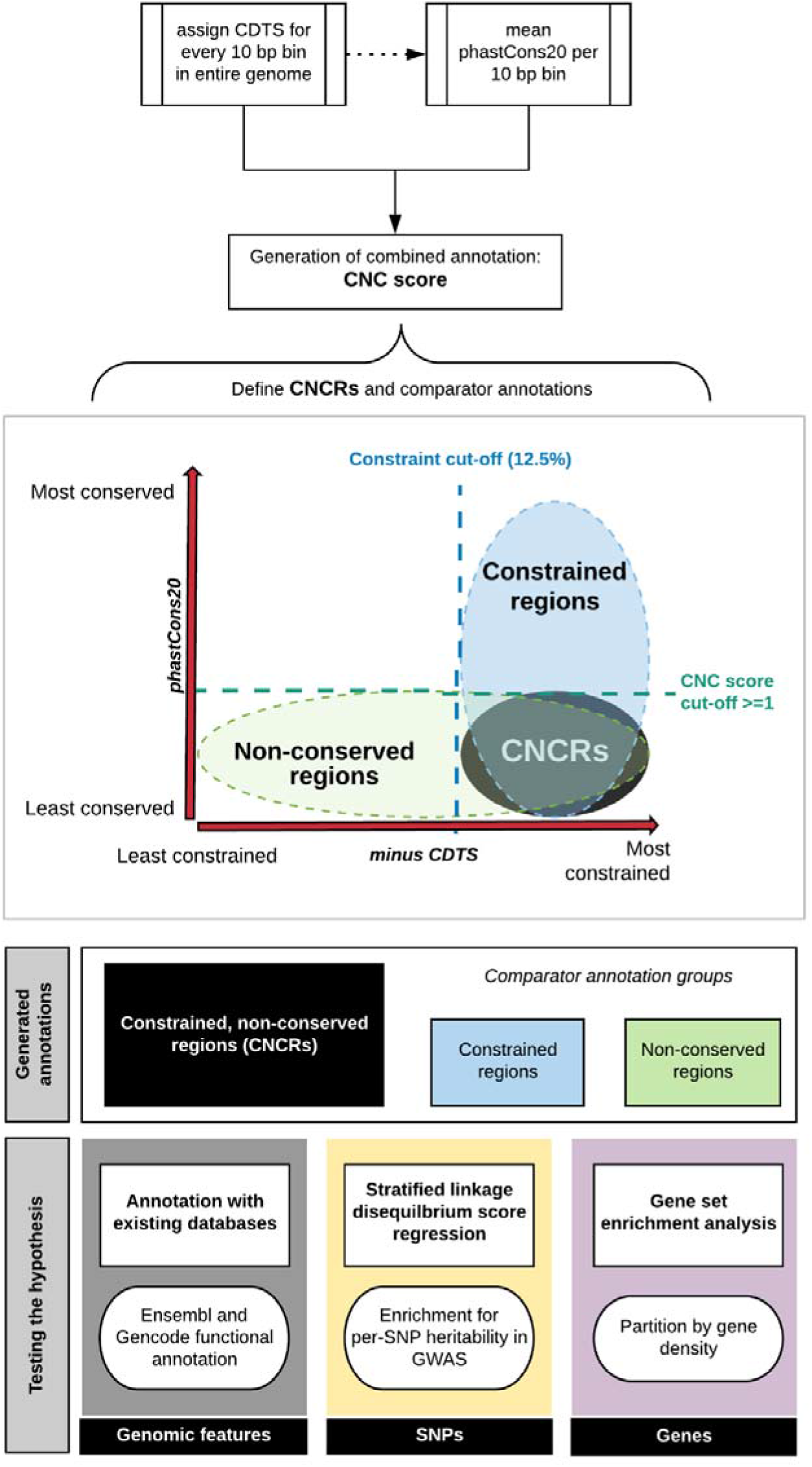
Workflow of study and schematic demonstration of annotation groups. The workflow depicts the processes involved in creation of the annotation with set parameters for each of the three groups of annotations generated and the processes involved in hypothesis-testing. CNC scores = constrained, non-conserved scores; CNCRs = constrained, non-conserved regions, CDTS = contextdependent tolerance score. Minus CDTS score is used as a lower score of CDTS corresponds to a more constrained region.

### Investigating the relationship between CNCRs and existing annotation

To investigate the relationship between CNC scores for genomic regions and genomic features, we calculated the distribution of CNC scores across genomic features defined by GENCODE v.53^19^ and Ensembl v.92^20^. We restricted our analysis to the 12.5% most constrained regions only (31,115,616 ten bp bins) and segregated these regions into equally-sized deciles ranked on the basis of CNC scores such that the highest decile (90 – 100 decile) represented a high CNC score containing the most constrained and least conserved sequences. Each 10bp region was then assigned a single overlapping genomic feature. To avoid conflicts arising from overlapping GENCODE and Ensembl definitions, we preferentially assigned a single genomic feature to a given region by prioritising features^11^ as described in **Supplementary Table 1**.

### Enrichment of common-SNP heritability in brain-related phenotypes for CNCRs

Stratified linkage disequilibrium score regression (LDSC) was used to assess the enrichment of common-SNP heritability for a range of complex diseases and traits within our annotation^21,22^. Stratified LDSC makes use of the increased likelihood of a causal relationship in a block of SNPs in linkage disequilibrium (LD) to correct for confounding biases that include cryptic relatedness and population stratification in a polygenic trait^22^. Using established protocols (https://github.com/bulik/ldsc/wiki), we tested whether SNPs located within our annotation contributed significantly to SNP-heritability after controlling for a range of other annotations described within the baseline mode (v.1.2). This analysis generates a coefficient z-score, from which we calculated a one-tailed coefficient p-value. Stratified LDSC regression analyses were also run to incorporate background SNPs defined as all SNPs in the genome that include a CDTS and phastCons20 annotation, to avoid over-estimation of the contribution to SNP-heritability. We assessed the annotation for SNP-heritability enrichment in complex brain-related disorders and phenotypes of intelligence^23^, Alzheimer’s disease^24^, Parkinson’s disease (excluding 23&Me participants)^25^, schizophrenia^26^ and major depressive disorder (excluding 23&Me participants)^27^ (**Supplementary Table 2**). We considered SNPs within CNCRs and its two constituent groups (**Figure 1**) which fall either into constrained only or non-conserved only annotations as defined respectively by: (i) CNCRs annotation: SNPs with a given CNCR density; (ii) Constrained annotation: SNPs located within the 12.5% most constrained regions of the genome irrespective of conservation score; (iii) Non-conserved annotation: SNPs located within relatively non-conserved genomic regions with a conservation rank determined by the rank of the first quartile phastCons20 score at a CNC score of 1 (rank ≤ 25,623,592) (irrespective of constraint score). We provide Bonferroni-corrected *p*-values, which account for the number of annotation categories and GWASs tested (total of 15 conditions).

### Generation of a gene-based metric for CNCRs and gene set enrichment analysis

To generate a metric of human-specific constraint, which could be applied to a gene rather than a 10bp region, we calculated the density of CNCRs within each gene, the length of which was defined by the transcription start and stop sites for that gene (GRCh38.v97). We used g:ProfileR (R Package)^28^ for gene set enrichment analysis. We used the three sets of tested annotations incorporating genes that fell into CNCRs, constrained regions and non-conserved regions in the gene set enrichment analysis as previously described for LDSC annotation and as defined in **Figure 1**. The background gene list in all analyses comprised 49,644 genes from all regions of the genome with a CDTS and phastCons20 annotation. The correction method was set to g:SCS to account for multiple testing^28^. We used REViGO^29^ to summarise the significant GO terms, and to derive the term frequency, which is a measure of GO term specificity.

To further characterise CNCR density within genes associated with disease, we first studied phenotype relationships of all Mendelian genes within the Online Mendelian Inheritance in Man (OMIM) catalogue (http://api.omim.org)^30^. We compared the CNCR density of all neurologically-relevant OMIM genes to all genes within CNCR annotation. Secondly, in order to investigate the CNCR density within genes associated with complex disorders, we used the Systematic Target OPportunity assessment by Genetic Association Predictions (STOPGAP) database, a catalogue of human genetic associations mapped to effector gene candidates derived from 4,684 GWASs^31^. We selected for genes associated with SNPs that surpassed a genome-wide significant p-value of 5×10^−8^ and which fulfilled medical subject heading for associated neurological/behavioural diseases. We used these sets to identify potential genes of interest associated with brain-related disorders which carry a high CNCR density.

### Sequencing of *APOE* transcripts in human brain

Focussing on a human-specific event identified within *APOE* from the preceding analyses, we used Sanger sequencing of cDNA reverse transcribed from pooled human hippocampus poly-A-selected RNA (Takara/Clontech 636165) to support the presence of the human-specific intron-3 retention event identified within *APOE* (GRCh38: chr19:44907952-44908531). For the reverse transcription, we used 500 ng of input RNA, with 10 mM dNTPs (NEB N0447S), VN primers and strand-switching primers (Oxford Nanopore Technologies SQK-DCS109), 40 units of RNaseOUT inhibitor (Life Technologies 10777019) and 200 units of Maxima H Minus reverse transcriptase with 5X reverse transcription buffer (ThermoFisher EP0751). PCR amplification of the cDNA was performed using primer pairs designed to span across intron-3 and exon 4 (P2-4) and intron-3 alone (P5) of *APOE* (ENST00000252486.9) (**Supplementary Table 3**). PCR was performed using Taq DNA polymerase with Q-solution (Qiagen) and enzymatic clean-up of PCR products was performed using Exonuclease I (ThermoScientific) and FastAP thermosensitive alkaline phosphatase (ThermoScientific). Sanger sequencing was performed using the BigDye terminator kit (Applied Biosystems) and sequence reactions were run on ABI PRISM 3730xl sequencing apparatus (Applied Biosystems). Electropherograms were viewed and sequences were exported using Sequencher 5.4.6 (Gene Codes). Sequences were aligned against the human genome (hg38) using BLAT and visually inspected for confirmation of validation.

### Analysis of public RNA-sequencing data

We used publicly-available short read RNA-sequencing data from human brain post-mortem samples provided by Genotype-Tissue Expression Consortium (GTEx) v.7.1^32^ and the Religious Orders Study and Memory and Aging Project (ROSMAP) Study^33^ and to quantify the human-specific intron-3 retention event in *APOE* highlighted by our analysis. For GTEx data, we used pre-aligned files available from recount2 (https://jhubiostatistics.shinyapps.io/recount/)^34^. Both studies within ROSMAP are longitudinal clinicopathological cohort studies of aging and/or Alzheimer’s disease. We downloaded BAM files for ROSMAP bulk-RNA sequencing data from the Synapse repository (https://www.synapse.org/#!Synapse:syn4164376) for analysis. To quantify the intron-3 retention event, we calculated the coverage of intron-3 expression normalised for the coverage across the entire *APOE* gene, as defined by the transcription start and end sites. To quantify splicing of intron-3, we calculated the number of exon-3 to exon-4 junction reads (defined as reads mapping with a gapped alignment), normalised for all *APOE* junction reads detected and currently within annotation. We used a ratio of the normalised coverage to normalised junction count over intron-3 as an estimate of the proportional use of the intron-3-retaining transcript, such that a high ratio is associated with a higher usage of intron retention within both GTEx and ROSMAP data. Based on existing ROSMAP results^35^ and principal component analysis of fragments per kilobase million (FKPM) data, we incorporated covariates to account for the effect of batch, RNA integrity number (RIN), postmortem interval (PMI), study index, ethnicity, age at death and sex on estimates of intron-3-retaining transcript usage. Using the resulting mixed linear model, we compared the intron-3 retention normalised coverage to junction ratio across clinical disease states, pathological states and *APOE* status in 634 post-mortem brain samples.

### Data availability

We release our annotation of CNC score as an interactive visualisable track via online platform vizER and provide a publicly-downloadable table of CNCR density for genes within our annotation (https://snca.atica.um.es/browser/app/vizER).

## RESULTS

### Genomic regions with high constraint, but not conservation were enriched for regulatory, non-coding genomic features

CNC scores, which combine information from CDTS and phastCons20, were used to capture evidence of disparity between constraint and conservation within a genomic region (**Figure 1**). We investigated the relationship between CNC scores and known genomic features within the most constrained portion of the genome (top 12.5%). This analysis demonstrated clear patterns of enhancement and depletion for genomic elements across CNC scores, which significantly differed from similar analyses performed using constraint metrics alone^11^ (**Figure 2a**). Among constrained genomic regions with the highest CNC scores (90-100 decile, signifying high constraint, but low conservation) we saw a depletion for coding elements of 27-fold relative to genomic regions with the lowest CNC scores. This contrasts with the pattern using constraint metrics alone where the most constrained genomic regions are highly enriched for coding exons^11^. On the other hand, promoter, promoter-flanking, and non-coding RNA features were over-represented in the highest compared to the lowest CNC deciles by 4.7, 1.9 and 1.5-fold respectively. Thus, genomic regions with high CNC scores are enriched for regulatory, non-coding genomic features.

**Figure 2.**
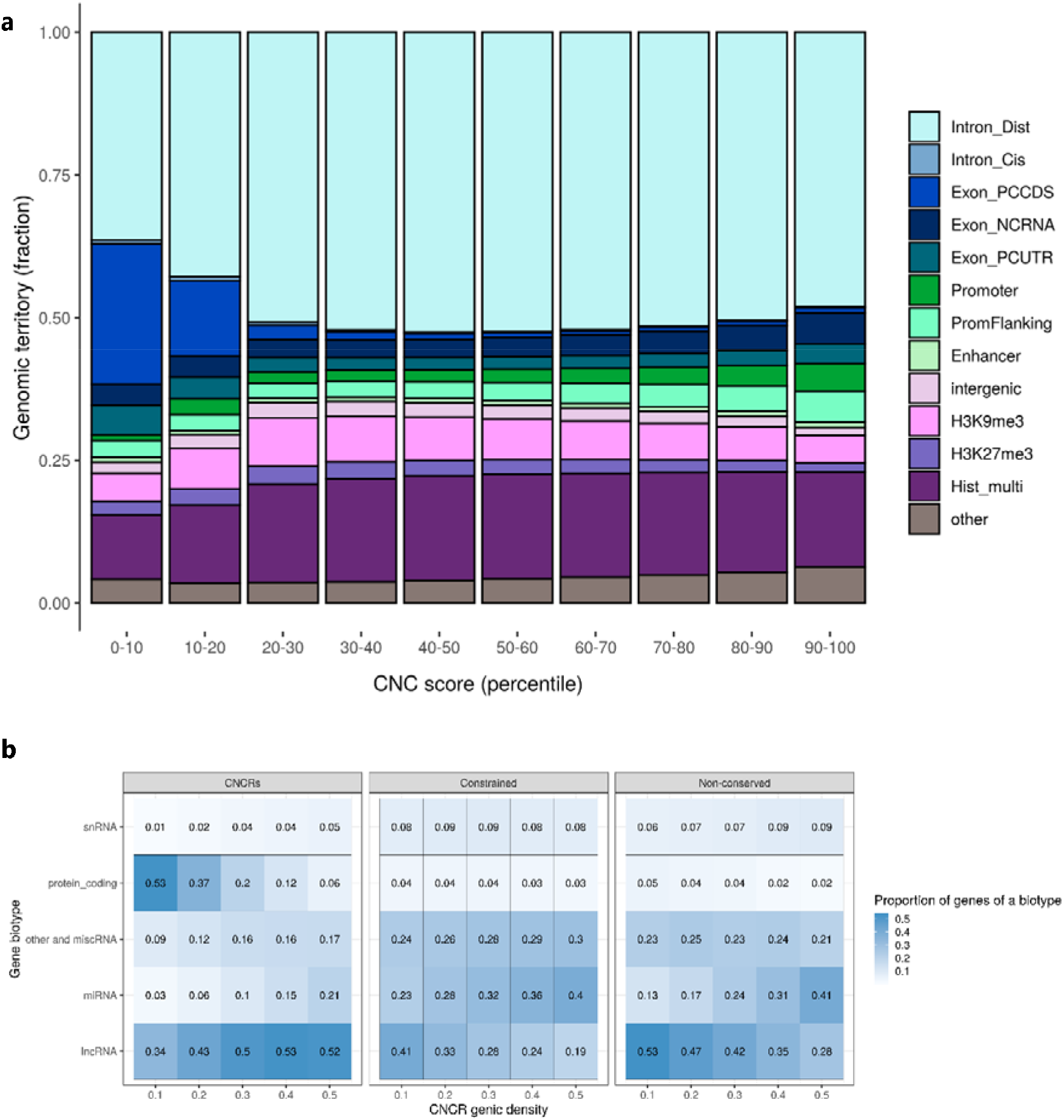
Composition of the constrained genome, partitioned by constrained, non-conserved (CNC) scores (a) and proportion of biotypes of genes in our annotation (CNCRs) and in the comparator annotations (constrained and non-conserved regions) (b). The description for each genomic feature is shown in Supplementary Table 1. The barplot in Panel **a** shows the genomic features for the 12.5% most constrained regions with CNC scores portioned by decile, such that the highest decile (90 – 100) represents the most constrained and least conserved regions. Description of gene biotypes in Panel **b** are taken from Ensembl^20^. The heatmap demonstrates the proportion of genes of a certain biotype within the three separate annotations within each genic CNCR density cutoff. Protein coding is defined by a gene that contains an open reading frame. The subclassified components of long non-coding RNA (lncRNA) found in the annotations are: Antisense – has transcripts that overlap the genomic span (i.e. exon or introns) of a protein-coding locus on the opposite strand; lincRNA (long interspersed ncRNA) – has transcripts that are long intergenic noncoding RNA locus with a length >200bp; non-coding RNA is further subclassified into miRNA (microRNA); siRNA (small interfering RNA); snRNA (small nuclear RNA) and miscellaneous RNA (includes snoRNA (small nucleolar RNA), tRNA (transfer RNA)). Pseudogenes are similar to known proteins but contain a frameshift and/or stop codon(s) which disrupts the open reading frame. These can be classified into processed pseudogene – a pseudogene that lacks introns and is thought to arise from reverse transcription of mRNA followed by reinsertion of DNA into the genome and unprocessed pseudogene – a pseudogene that can contain introns since produced by gene duplication.

### Genes with the highest density of CNCRs are enriched for long non-coding RNA

Next, we applied a CNC score cut-off of ≥ 1 (signifying a two-fold higher ranking in constraint than conservation) to define a set of genomic regions which were constrained, but not conserved (termed CNCRs). Next, we wanted to investigate whether CNCRs could be used to identify specific genes of interest. With this in mind, we used CNCR density to identify gene sets which might be expected to contribute most to human-specific phenotypes. Consistent with the findings above, we found that as the CNCR density threshold was increased to define the gene sets of interest, there was a marked reduction in the proportion of protein-coding genes (β-coefficient between proportion and CNCR density = −1.061 and false discovery rate (FDR)-corrected p = 0.00162), and an increase in the proportion of long non-coding RNA (lncRNA, β-coefficient 0.385 and FDR-corrected p = 0.0161), and microRNA-encoding genes (miRNA, β-coefficient 0.394 and FDR-corrected p = 0.00116) (**Figure 2b**). Interestingly, this relationship was not clearly observed when considering unprocessed snRNA and other RNAs (**Figure 2b**). In order to determine whether the relationship between CNCR density and gene biotype was driven by sequence constraint or conservation, we also generated comparator gene lists based on constrained-only and non-conserved regions alone. Importantly, lncRNA and protein-coding gene proportions do not follow the same directionality with increasing density when constraint or non-conservation alone is considered (**Figure 2b**). Thus, this analysis highlighted the specific importance of lncRNAs as compared to other classes of non-coding RNAs in driving human-specific patterns of gene expression.

### Significant enrichment of heritability for neurologically-relevant phenotypes

Given the enrichment of regulatory features within genomic regions with a high CNC score, we postulated that such regions could also be enriched for disease risk. In order to study this, we investigated CNCRs for evidence of enriched heritability for a range of complex neurologically-relevant phenotypes (**Supplementary Table 4**). After Bonferroni correction for multiple testing, we found that CNCRs exhibited significant enrichment in heritability for intelligence (coefficient p = 4.19×10^−24^); Parkinson’s disease (coefficient p = 4.65×10^−5^); major depressive disorder (coefficient p = 2.95×10^−8^) and schizophrenia (coefficient p = 5.26×10^−19^), but not for Alzheimer’s disease (**Figure 3**). While a significant enrichment in heritability for intelligence, major depressive disorder and schizophrenia were also observed in the constrained regions alone (and to a lesser extent, nonconserved regions), we note that the regression coefficient for CNCRs was at least two-fold larger for the CNCR annotation compared to the constrained annotation (**Supplementary Table 4**). Similarly, significant enrichment in heritability for Parkinson’s disease was only observed in CNCRs. Thus, by combining metrics for both constraint and conservation in our annotation, we derived an independent annotation that shows a higher level of enrichment in heritability for neurologically-related phenotypes than annotations based on constraint or conservation alone.

**Figure 3.**
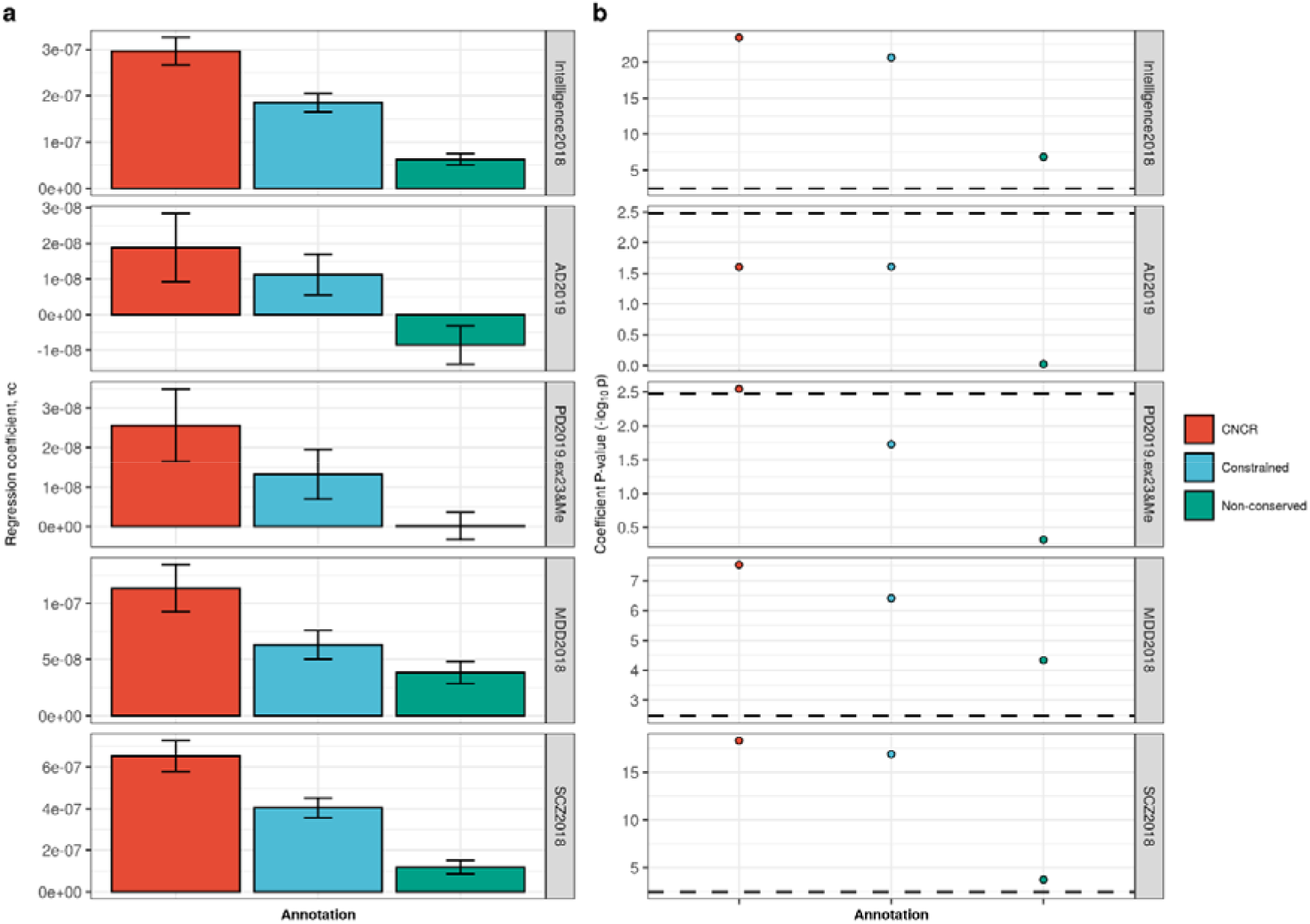
Stratified-LDSC analysis across five traits comparing CNCRs with its constituent constrained and non-conserved annotations. Panel **a** shows the regression coefficient. Panel **b** shows the regression coefficient −log(p-value) with the dotted line showing the Bonferroni-corrected p-value of 0.00333. GWASs were as follows: Intelligence2019 = intelligence GWAS, AD2018 = Alzheimer’s disease GWAS, PD2019.ex23&Me = Parkinson’s disease GWAS without 23 and Me data, MDD2018 = Major depressive disorders GWAS and SCZ2018 = schizophrenia GWAS (**Supplementary Table 2**).

### The proportion of enriched gene sets with neurologically-related GO terms increases in genes with the highest density of CNCRs

To investigate these findings further, we defined gene sets based on their CNCR density and analysed their GO term enrichment. We assessed gene sets defined across a range of CNCR densities (> 0.0 to ≥ 0.5 at 0.1 increments). We found that the proportion of neurologically-associated GO terms with significant enrichments (g:SCS-corrected p < 0.05) increased among gene sets with increasing CNCR gene densities (**Supplementary Figure 2**). Importantly, a similar analysis of gene sets defined by constraint alone or non-conservation alone did not contain any neurologically-enriched GO terms (**Figure 4**). We identified the gene set with the highest proportion of nervous system-related terms at a CNCR genic density of 0.3 (**Supplementary Figure 2**). The only GO terms specific to a tissue process were related to the nervous system (**Figure 4, Supplementary Table 5**) and spanned terms such as neuronal development (GO:0048663, corrected p = 5.46×10^−7^) and spinal cord differentiation (GO:0021515, corrected p = 3.64×10^−7^). The remaining significantly enriched GO terms related to ubiquitous processes including protein targeting (GO:0045047, p = 9.93×10^−4^) and DNA binding (GO:0043565, p = 4.81×10^−4^). Of note, analysis of gene sets defined on the basis of constraint alone revealed no enrichment of neurologically-associated terms, but instead significant enrichment of vascular system-related GO terms (GO:0048514 blood vessel morphogenesis, corrected p = 3.96×10^−37^ and GO:0072358 cardiovascular system development, p = 8.53×10^−36^). As might be expected based on the rapid and potentially divergent evolutionary pressures, the analysis of gene sets defined on the basis of non-conservation alone demonstrated the significant enrichment of immune and skin-related GO terms (GO:0002250 adaptive immune response, p = 4.02×10^−10^ and GO:0043588 skin development, p = 2.33×10^−4^). Taken together, these results demonstrate that using CNCR density, genes important in nervous system development and implicated in neurological disease can be identified.

**Figure 4.**
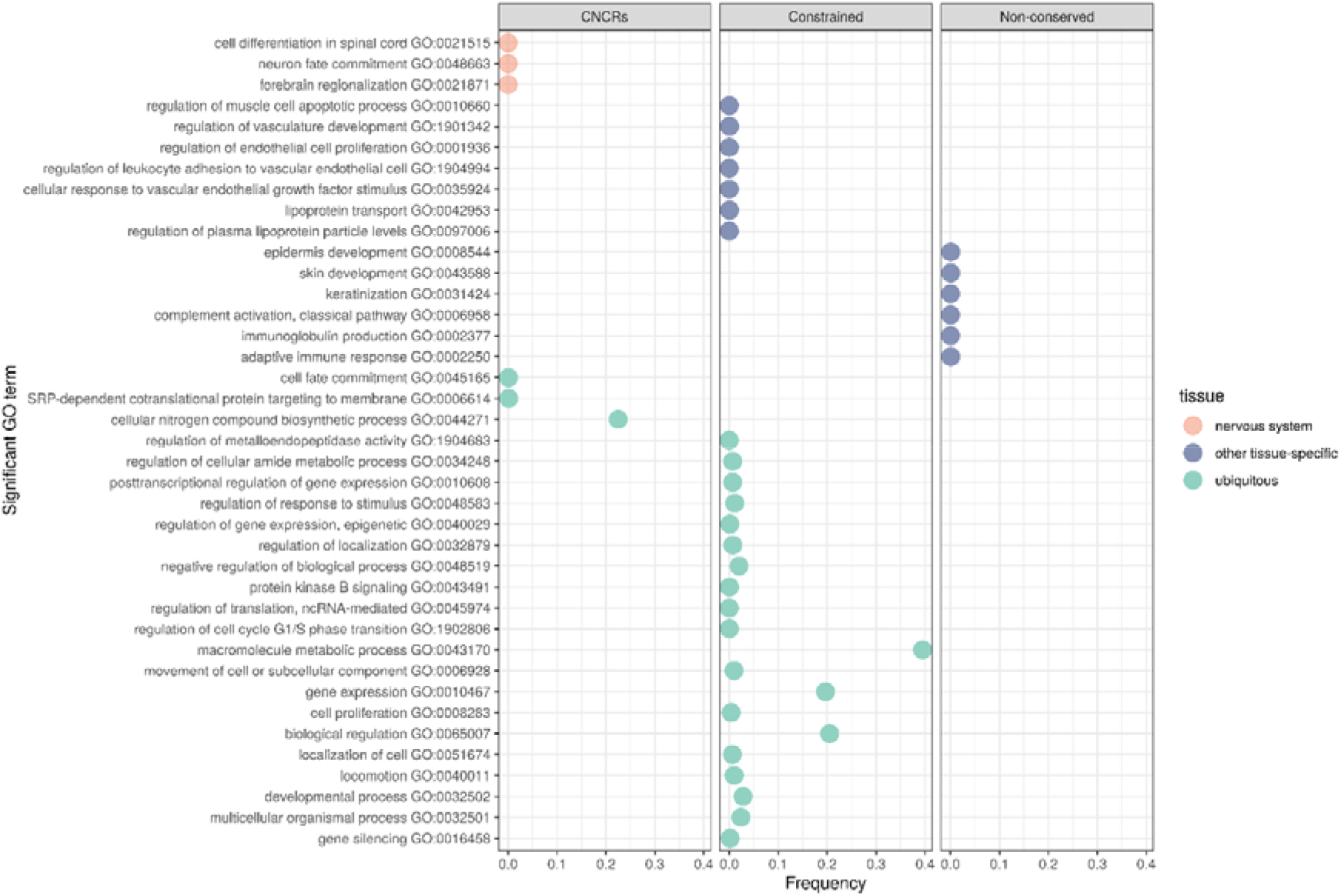
Summarised enriched gene sets for terms specific for neurological gene sets, other tissues and all tissues (non-neurological) as defined by Gene Ontology (GO). Plot comparing annotation of interest (CNCRs) and comparator annotations which only use constraint or nonconserved metrics. Frequency, derived from REViGO^29^, the percentage of human proteins in UniProt which were annotated with a GO term, i.e. a higher frequency denotes a more general term.

### CNCR annotation highlights an intron-3 retaining transcript of *APOE*

Next, we investigated the distribution of CNCR density across Mendelian genes associated with a neurological phenotypes (as defined within OMIM^30^) and genes implicated in complex brain-relevant phenotypes (as defined within STOPGAP^31^). We noted that the median CNCR density was significantly higher in OMIM genes with a neurological phenotype compared to all other genes (median CNCR density of neurological OMIM genes = 0.0924, IQR = 0.0567 – 0.143; median CNCR density of all other genes = 0.083, IQR = 0.043 – 0.153; Wilcoxon rank sum test p = 1.8×10^−6^). While genes associated with complex brain-relevant phenotypes did not have a significantly higher CNCR density when compared to all other genes, we still identified 31 genes with a CNCR density of greater than 0.2 and seven genes with a CNCR density of greater than 0.3 (*APOE, PHOX2B, SSTR1, HCFC1, HAPLN4, CENPM* and *IQCF5*). Of these genes, *APOE* had the highest CNCR density with a value of 0.552.

Given the high CNCR density of *APOE*, its importance as a disease locus for Alzheimer’s disease and other neurodegenerative diseases^36^ and the long-standing interest in the lineage specificity of *APOE^8,37^* (specifically the differences in the □4 allele between humans and non-human primates^1^), we chose to focus on this gene. We tested whether intragenic analysis of *APOE* could identify specific exons or transcripts of interest. We compared CNCR density, constraint and conservation scores across the length of the gene showing that CNCRs provide a highly granular annotation (**Figure 5**). Using this approach, we identified a region of high CNCR density within intron-3 of *APOE*. Although no intron-3 retaining transcript is currently annotated in Refseq and Ensembl, an intron-3 retention event has previously been reported and implicated in the regulation of *APOE* expression^38,39^. To validate the existence of this transcript, we performed Sanger sequencing of polyA-selected RNA derived from human hippocampal tissue. This demonstrated that no recursive splicing occurred as the full-length intron-3 sequence was retained and flanked by both exon-3 and exon-4 (**Supplementary Figure 3**).

**Figure 5.**
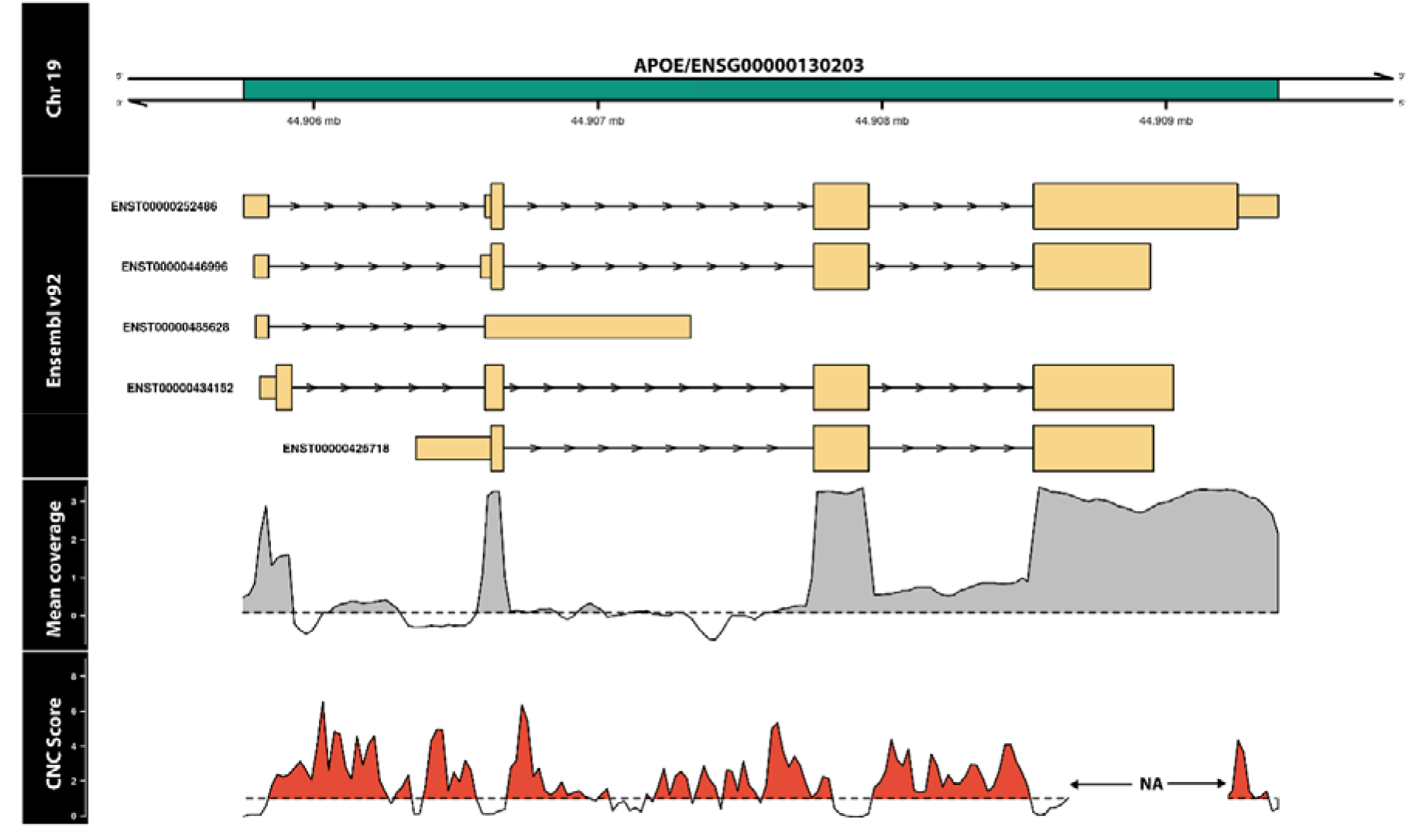
Annotation with CNCRs is highly granular and shows *APOE* to have a high density of CNCRs throughout its length especially in association with an intron-3 retention event in the human hippocampus. The first track represents the genomic location of *APOE* within Chromosome 19. The second track shows the known transcripts, currently within annotation in Ensembl v.92. The mean coverage (log10 scale) in the hippocampus shown here is greater than zero (denoted by the grey shaded area) across intron-3 highlighting a potential novel expressed region. In the last track, CNC scores above the black dashed line and shaded in red fulfil criteria for a CNCR.

In order to obtain further insights into the biological significance of the intron-3 retaining *APOE* transcript, we leveraged publicly-available RNA-sequencing data covering 11 regions of the human central nervous system provided by the GTEx v.7^32^. Using an annotation-independent approach to identify genomic regions producing stable transcripts^40,41^, we identified a region of significant expression encompassing intron-3 of *APOE* and the flanking coding exons in all brain tissues (**Figure 6a**). These data not only support the existence of an intron-3 retaining *APOE* transcript that is not entirely attributable to pre-mRNA transcripts or driven by background noise in sequencing, but also provide a means of estimating its usage across the human brain.

**Figure 6.**
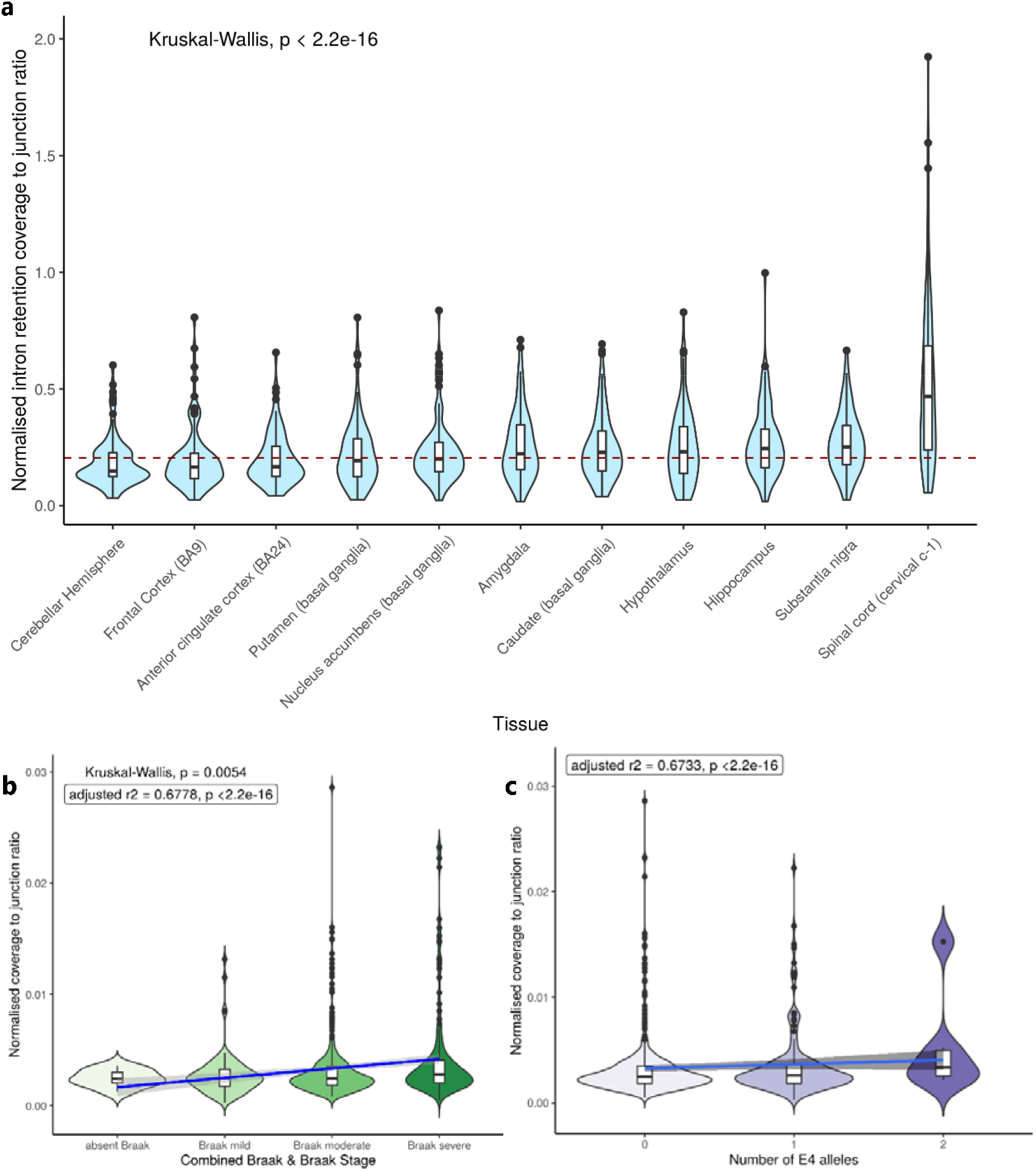
Quantification of intron retention usage by its normalised coverage to junction ratio across brain tissues within GTEx (a). Normalised coverage to junction ratio of the *APOE* intron-3 retention event in bulk RNA sequencing data of post-mortem dorsolateral prefrontal cortex tissue samples from 634 individuals recruited within ROSMAP studies across Braak and Braak staging (b) and *APOE* □4 allele status (c). In Panel **a**: red dashed horizontal line presents the median normalised intron retention coverage to junction ratio within central nervous system tissues in GTEx. Number of samples within each of the tissue groups were as follows: amygdala – 72; anterior cingulate cortex – 84; caudate – 117, cerebellar hemisphere – 105; frontal cortex – 108; hippocampus – 94; hypothalamus – 96; nucleus accumbens – 113; putamen – 97; spinal cord – 71; substantia nigra – 63. In panels **b** and **c**, the blue line represents the linear regression fit with the grey shaded area representing +/- 95% confidence interval. Braak and Braak staging is a measure of severity of neurofibrillary tangle based on location. To improve the power of the study, we merged Braak and Braak stages I and II to “Braak mild stage”, Braak and Braak stages III and IV to “Braak moderate” and Braak and Braak stages V and VI to indicate “Braak severe” stage. For number of *APOE* □4 alleles, a heterozygous state is represented by “1” and homozygous state by “2”.

Thus, in order to compare usage of this transcript across different CNS regions, we calculated the ratio of normalised intron-3 expression (a measure of intron-3 retaining transcripts) to the normalised expression of exon-3/exon-4 spanning reads (a measure of transcripts splicing out intron-3). We see that there is evidence of the usage of the intron-3 retaining *APOE* transcript in all central nervous system regions from GTEx data (**Figure 6a**). However, there are also significant differences among brain regions (Kruskal-Wallis p < 2.2e^−16^) with the usage of the intron-3 retaining eventconserved and thus evolutionarily being highest in the spinal cord, substantia nigra and hippocampus (**Figure 6a**).

In summary, we confirmed the existence of an unannotated human-specific non-coding transcript of *APOE* and identified differential usage of this transcript across the human brain. In this way, we demonstrate the utility of combining CNC scores with transcriptomic data, which we have made easier though the addition of a CNC score track within the platform vizER (https://snca.atica.um.es/browser/app/vizER).

### Usage of the intron-3 retaining transcript of *APOE* correlates with Alzheimer’s disease pathology and *APOE* genotype

We noted that among the brain tissues with the highest usage of the intron-3 retaining transcript of *APOE* are those that show selective vulnerability for neurodegeneration, namely the hippocampus in the context of Alzheimer’s disease, the substantia nigra in the context of Parkinson’s disease and the spinal cord in the context of amyotrophic lateral sclerosis. Given that *APOE* is one of the most important genetic risk factors for Alzheimer’s disease, we leveraged publicly-available RNA-sequencing data from the ROSMAP studies to quantify the usage of the intron-3 retaining transcript of *APOE* in post-mortem dorsolateral prefrontal cortex brain tissue derived from individuals with Alzheimer’s disease (n = 222), mild cognitive impairment (MCI) (n = 158) compared to control individuals (defined as the final clinical diagnosis blinded to pathological findings, n = 202). We found that the proportion of the intron-3-retaining transcript was higher (p < 2.2e^−16^) in dorsolateral prefrontal cortex tissue from individuals with clinically-diagnosed Alzheimer’s disease and MCI patients versus control participants. Partitioning this further on the basis of pathology, we see an increase in intron-3 retaining transcript usage with more severe Braak and Braak pathology for neurofibrillary tangles (adjusted r^2^ 0.678, p < 2.2e^−16^) (**Figure 6b**). Consistent with these findings, we also found a significant increase in transcript usage with lower CERAD stage, indicating higher amyloid plaque pathology (adjusted r^2^ 0.673, p < 2.2e^−16^). Finally, we investigated the relationship between presence of the ε4 allele in *APOE* and usage of the intron-3 retaining transcript. We found a significant positive correlation between ε4 allele load and the proportion of intron-3 retaining transcript (adjusted r^2^ 0.673, p < 2.2e^−16^) (**Figure 6c**). This association remained significant after partitioning *APOE*-ε4 status by disease and accounting for tau and amyloid burden, showing that this association is likely to be independent of disease state. Taken together, these findings suggest that usage of the intron-3 retaining transcript may be regulated by *APOE*-ε4 status and may be involved in mediating the effect of *APOE* genotype, supporting a role for the presence of this lncRNA in disease risk and progression.

## DISCUSSION

The core aim of this study was to test the hypothesis that capturing human lineage-specific regions of the genome could provide insights into neurological phenotypes and diseases in humans. We generate and use an annotation based on existing knowledge of sequence conservation and sequence constraint within humans, which we term CNCRs. We use this annotation to prioritise genomic regions, genes and transcripts based on a high density of human lineage-specific sequence as determined by our CNCR annotation. We demonstrate the utility of this approach by showing that: the genomic regions we identify are enriched for SNP-heritability for intelligence and brain-related disorders; the genes we identify are enriched for neurologically-relevant gene ontology terms and genes causing neurogenetic disorders; and the existence of an intron-3 retaining transcript of *APOE*, the usage of which is correlated with Alzheimer’s disease pathology and *APOE*-ε4 status.

A major finding of this study is that CNCRs are enriched for regulatory, non-coding genomic regions. This is consistent with analyses performed by Ward and Kellis^14^, and highlights the potential functional importance of *non*-conserved and thus evolutionarily-recent non-coding regions subject to constraint. Furthermore, these findings suggest that CNCRs could provide a means of prioritising and potentially aiding the assessment of non-coding variants, an area of significant interest given that 88% of GWAS-derived disease-associated variants reside in non-coding regions of the genome^42^. We found evidence to support this view through heritability analyses for intelligence, Parkinson’s disease, major depressive disorder and schizophrenia with SNP-heritability not only enriched within CNCRs, but to a greater extent than would be expected using either conservation or constraint annotations alone. Considering heritability for intelligence, this phenotype is already known to also be enriched within annotations of brain-specific tissue expression and among several regulatory biological gene sets^23^, including neurogenesis, central nervous system neuron differentiation and regulation of synapse structure or activity^42^. These findings support our hypothesis that CNCRs identify genomic regions of functional importance with relevance to human brain phenotypes.

Our analyses of CNCR density within genes are consistent with these findings, highlighting both noncoding genes and those implicated in neurologically-relevant processes and diseases. Interestingly, CNCR annotation specifically highlighted lncRNAs as opposed to other non-coding RNAs. In particular, we observed a proportional increase in lncRNA enrichment with higher genic CNCR density, which could not be replicated using measures of sequence constraint or conservation alone. This observation is in keeping with previous studies that have shown most lncRNAs are tissuespecific with the highest proportion being specific to brain^43^. Similarly, the enrichment for nervous system-related pathways within CNCRs, which is representative of recent purifying selection, is in keeping with the lowest proportion of positively-selected genes being present in brain tissues from previous studies of mammalian organ development^44^. We also find enrichment of spinal cord-associated genes that may relate to the uniquely human monosynaptic corticomotoneuronal pathways implicated in human-specific dexterity and digital motor control^45,46^, the disruption of which may lead to amyotrophic lateral sclerosis^47^.

We noted that *APOE* was among the genes with the highest CNCR density across the genome and carried the highest CNCR density of all genes implicated in complex brain-relevant phenotypes (defined within the STOPGAP database^31^). Given that genetic variation within this gene and specifically *APOE*-ε4 status is not only the principal genetic risk factor for Alzheimer’s disease^48^ but also associated with risk for other neurodegenerative disorders, stroke and reduced lifespan^36^, this finding provides evidence for the value of CNCR annotation. Furthermore, within *APOE*, the CNCR annotation highlighted an intron-3 retention event not currently within annotation but which has been previously reported^38,39^. Using Sanger sequencing of cDNA derived from control human hippocampal tissue, we confirm the presence of an intron-3 retaining *APOE* transcript. We estimate the usage of the transcript from short read RNA-sequencing data and find variable levels across different brain tissues within GTEx^32^ with the highest usage in the spinal cord, substantia nigra and hippocampus, reflecting the brain regions most susceptible to selective vulnerability in disease. Using human dorsolateral prefrontal cortex RNA-sequencing data, we find that the intron retention event is significantly more abundant in patients with Alzheimer’s disease than controls and in those with more severe Braak and Braak pathology and amyloid burden as characterised by CERAD pathology. Furthermore, we see a dosage-dependent increase in the intron retention event with the *APOE*-ε4 allele that is independent of disease status. We propose that this novel transcript may be a means of regulating *APOE* in a disease state.

Given that we use existing measures of constraint and conservation to identify CNCRs, this analysis is fundamentally limited by the quality of these data. While the constraint metrics we used were derived from high depth sequencing, this is still restricted given the relatively high number of private genetic variants we each carry. In addition, analysis was limited to the high-confidence regions covering approximately 84% of the genome, so a significant proportion remained unannotated with CDTS metrics^11^. Similarly, our study of the relationship between CNCRs and known genomic features is limited by the annotation quality in existing databases. We have endeavoured to overcome some of these problems by creating a more detailed annotation combining both GENCODE and Ensembl data. The SNP-heritability estimates using stratified-LDSC analysis are limited by the quality of LD information underpinning the heritability calculations^21^.

Despite these limitations, we have been able to demonstrate the utility of CNCRs specifically in the identification of functionally important non-coding regions of the genome, genes and transcripts. We find that CNCRs across all forms of analyses highlight the significance of human lineage-specific sequences in the central nervous system and in the context of neurological phenotypes and diseases. We release our annotation of CNC scores via the online platform vizER (https://snca.atica.um.es/browser/app/vizER). Thus, the CNCR annotation we generate has the potential to provide additional disease insights beyond those explored within this study and as we anticipate the release of increasing quantities of WGS data in humans, will only improve in quality and value.

## AUTHOR CONTRIBUTIONS

ZC and DZ generated the annotation and conducted further analyses. ZC and RHR performed LDSC analysis. ZC and EKG generated cDNA and completed Sanger sequencing. ZC conducted the RNA-sequencing data analyses of *APOE*. DZ and SGR developed the vizER platform for visualisation of CNC scores. KD’S, AFB and JV helped with further analyses of RNA-sequencing data. IPDGC contributed PD GWAS data summary statistics. MR conceived and supervised the study. JB, SGAT, HH and JH provided further guidance on technical aspects of the study. All authors contributed to the writing and reviewing of the manuscript.

## ACKNOWLEDGEMENTS

We are grateful to the participants in the Religious Order Study, the Memory and Aging Project. ZC and RHR were supported by grants from the Leonard Wolfson Foundation. JH was supported by the UK Dementia Research Institute which receives its funding from DRI Limited, funded by the UK Medical Research Council, Alzheimer’s Society and Alzheimer’s Research UK. JH has also been funded by the Medical Research Council (award MR/N026004/1), Wellcome Trust (award 202903/Z/16/Z), Dolby Family Fund, National Institute for Health Research University College London Hospitals Biomedical Research Centre.

## SUPPLEMENTAL DATA

### Supplementary Data Figure Titles and Legends

**Supplementary Figure 1.**
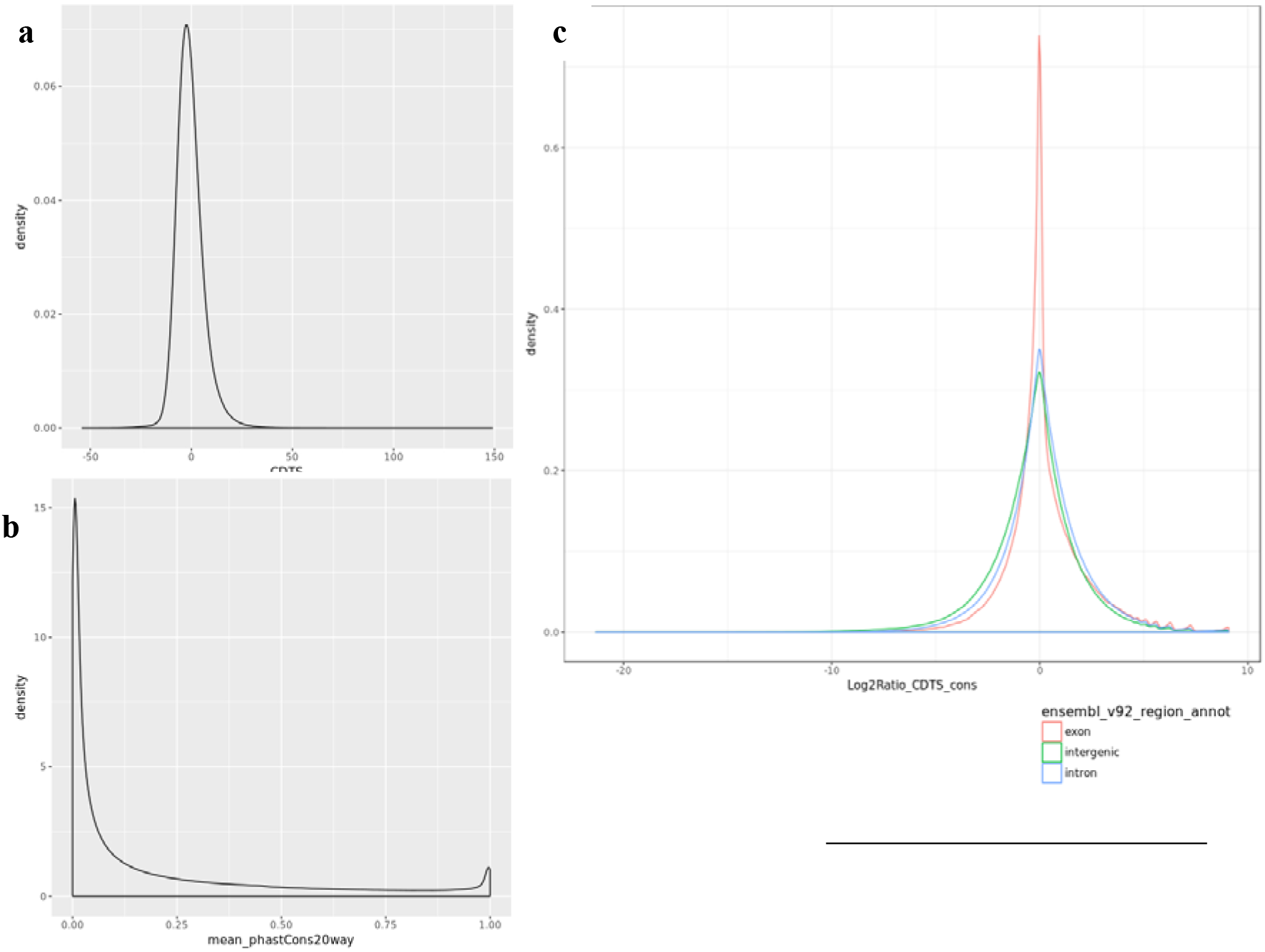
Kernal density plots of annotation metrics. Panel **a** depicts density plot of constraint (context dependent tolerance score (CDTS): a lower CDTS represents more constrained data). Panel **b** shows the density distribution of the mean phastCons20 scores per 10bp bin. Panel **c** shows the distribution of log2 ratio (CNC score), of the reverse ranked CDTS (so a higher rank pertains to higher constraint but lower CDTS) and ranked phastCons20 scores, partitioned by regions of exon, intron and intergenic as defined by Ensembl v.92.

**Supplementary Figure 2.**
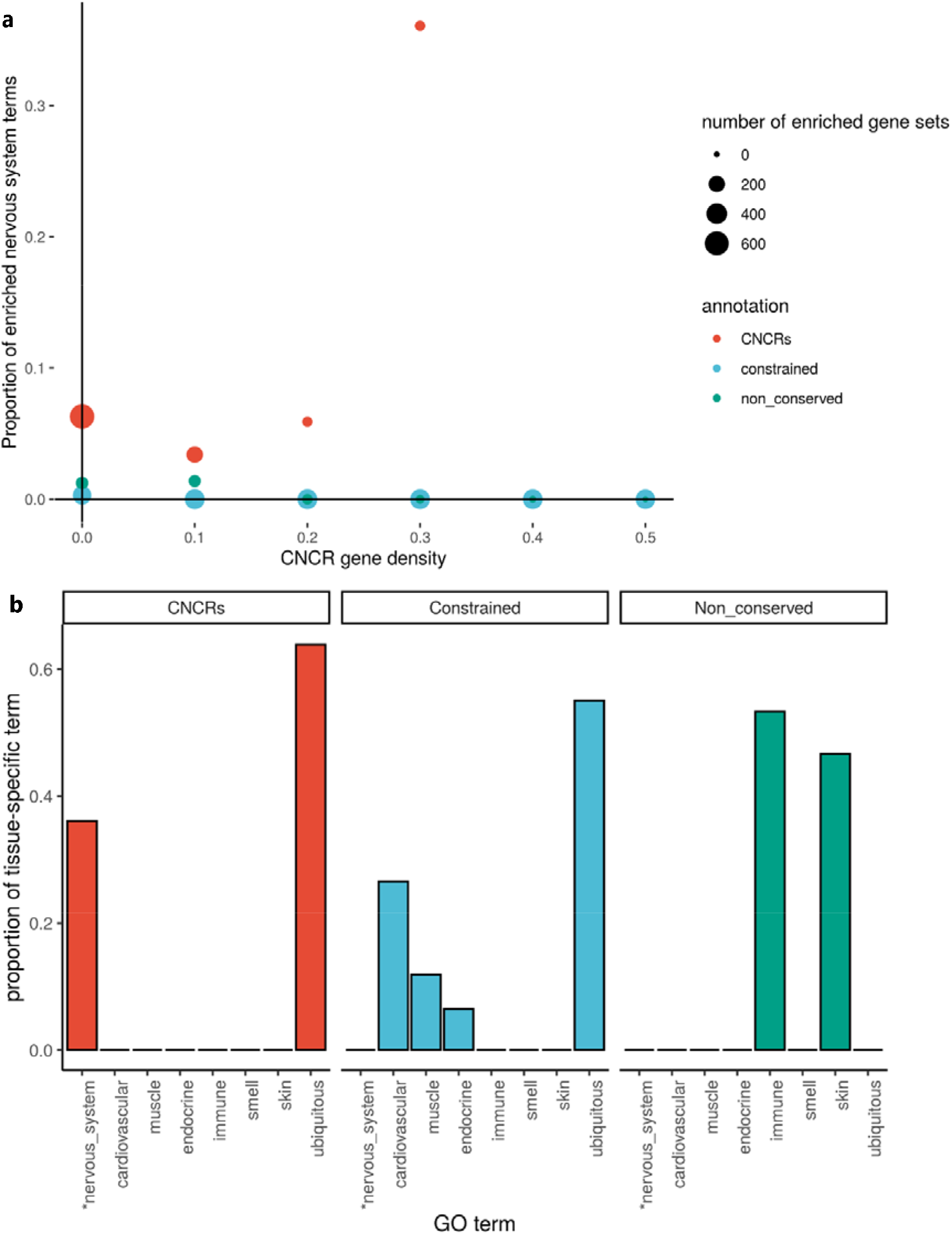
Proportion of enriched neurologically-related GO terms in the gene set analysis compared between the annotation of interest (CNCRs) and the comparator annotation sets (a). Proportion of neurologically-related GO terms at CNCR density of 0.3 and above (b).

**Supplementary Figure 3.**
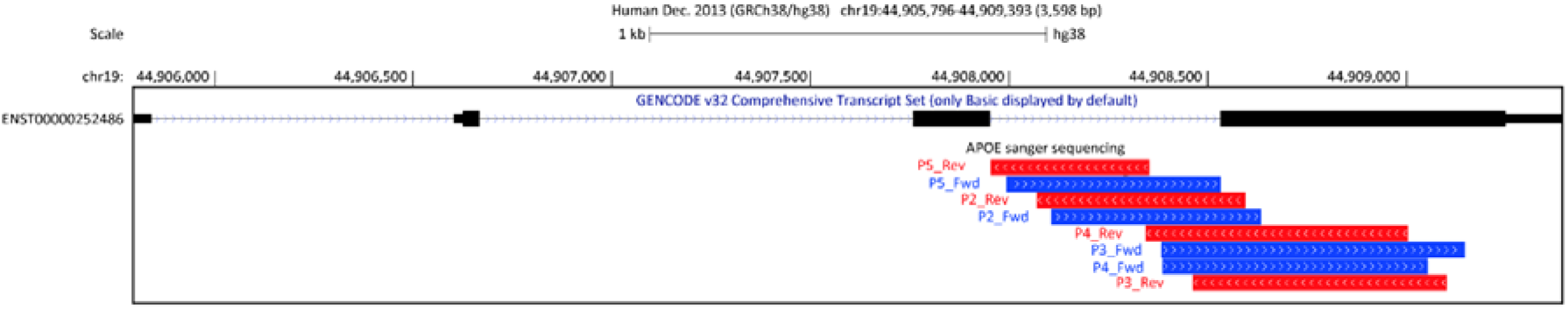
Sanger sequencing of human hippocampus cDNA using targeted primers within *APOE*, aligned to hg38. Primers as listed in Supplementary Table 3.

### Supplementary Tables

**Supplementary Table 1.** Annotation priority order for genomic feature. Genomic features are based on both Gencode and Ensembl. A priority order for annotation with a genomic feature is assigned to avoid conflict with overlapping features. The number of 10bp bins across the genome is also shown in the table.

| Annotation<br>priority order | Genomic feature | Number of<br>10bp bins | Description |
| --- | --- | --- | --- |
| 1 | Exon PCCDS | 1,453,269 | Exon, protein-coding sequence |
| 2 | Exon NCRNA | 1,156,726 | Exon, non-coding RNA, e.g. lincRNA |
| 3 | Exon PCUTR | 892,210 | Exon, protein-coding UTR |
| 4 | Promoter | 820,321 | Promoter |
| 5 | Promoter Flanking | 1,074,641 | Cluster with promoters or distal cis-regulatory elements |
| 6 | Enhancer | 251,636 | Enhancer |
| 7 | Intron, cis | 108,670 | Introns located in genes <10bp from splice-site |
| 8 | Intron, trans | 15,204,447 | Introns located in genes >10bp from splice-site |
| 9 | Intergenic | 689,419 | Not annotated in GenCode/ Ensembl |
| 10 | H3K9me3 | 2,082,553 | Only overlap with H3K9me3 |
| 11 | H3K27me3 | 777,409 | Only overlap with H3K27me3 |
| 12 | Multiple histones | 5,199,455 | Overlap with a combination of histone marks |
| 13 | Other | 1,404,860 | Includes open chromatin and unannotated features |

**Supplementary Table 2.** Genome-wide association studies used in the stratified LDSC analysis. The GWAS for Parkinson’s disease and major depressive disorder do not incorporate 23&Me data.

| **Disease** | **Author, Year, Reference** | **n case** |
| --- | --- | --- |
| Intelligence | Savage, 2018<sup>23</sup> | 269,858 |
| Alzheimer's disease (AD) | Jansen, 2018<sup>24</sup> | 71,880 |
| Parkinson's disease (PD) | Nalls, 2019 (excluding 23&Me data)<sup>25</sup> | 33,674 |
| Major depressive disorder (MDD) | Wray, 2018 (excluding 23&Me data)<sup>27</sup> | 59,851 |
| Schizophrenia (SCZ) | Pardiñas, 2018<sup>26</sup> | 40,675 |

**Supplementary Table 3.**
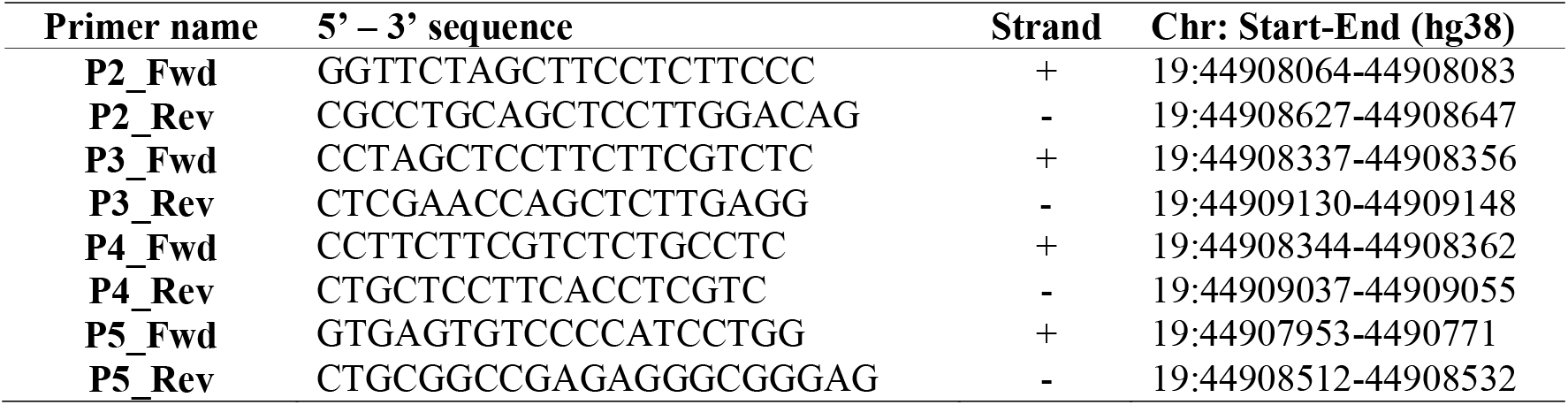
Primer positions and sequences used to validate the *APOE* intron-3 retention event.

**Supplementary Table 4.**
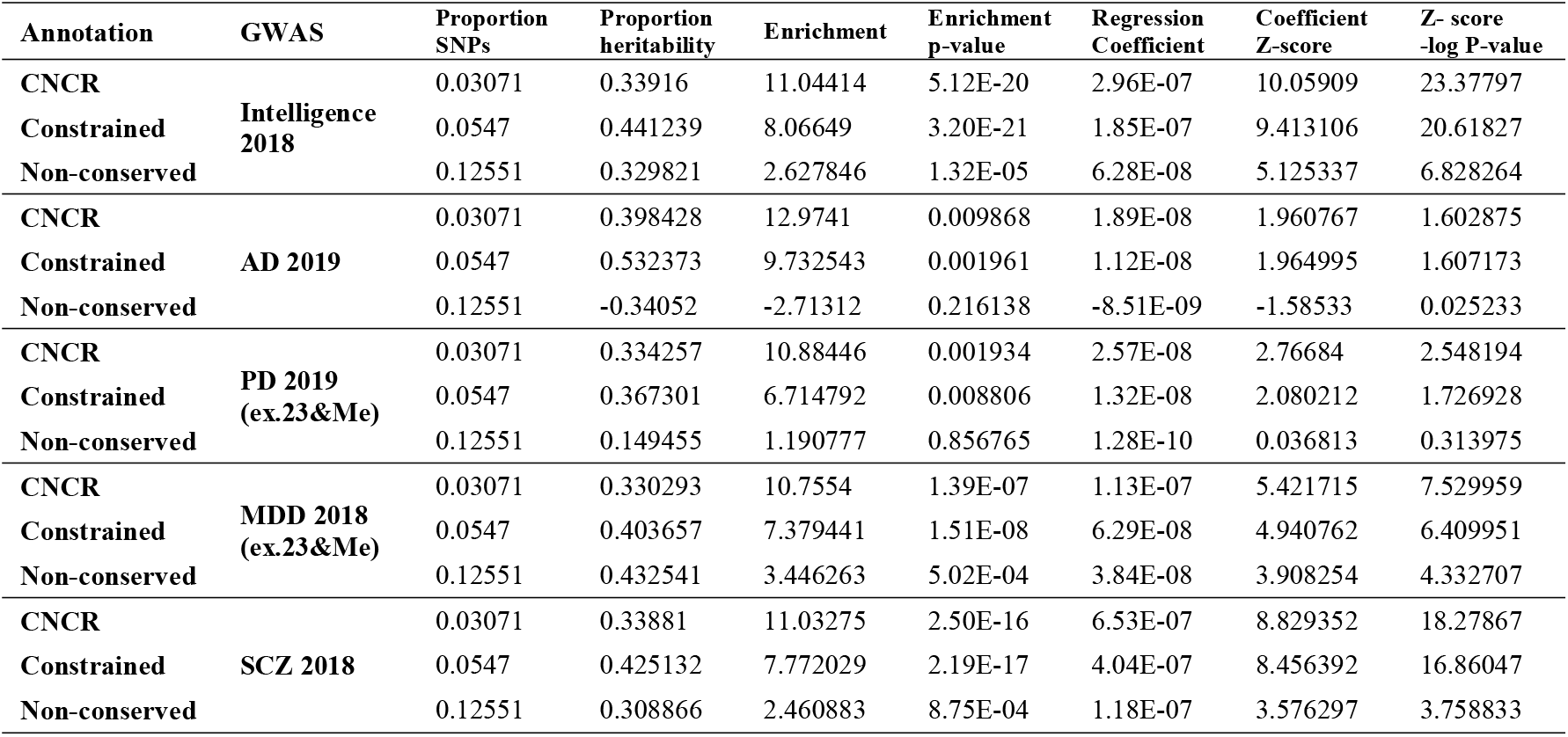
Results for heritability, enrichment, and regression coefficient from stratified LDSC analysis. The coefficient p-values are one-sided p-values calculated from the coefficient Z-score.

**Supplementary Table 5.**
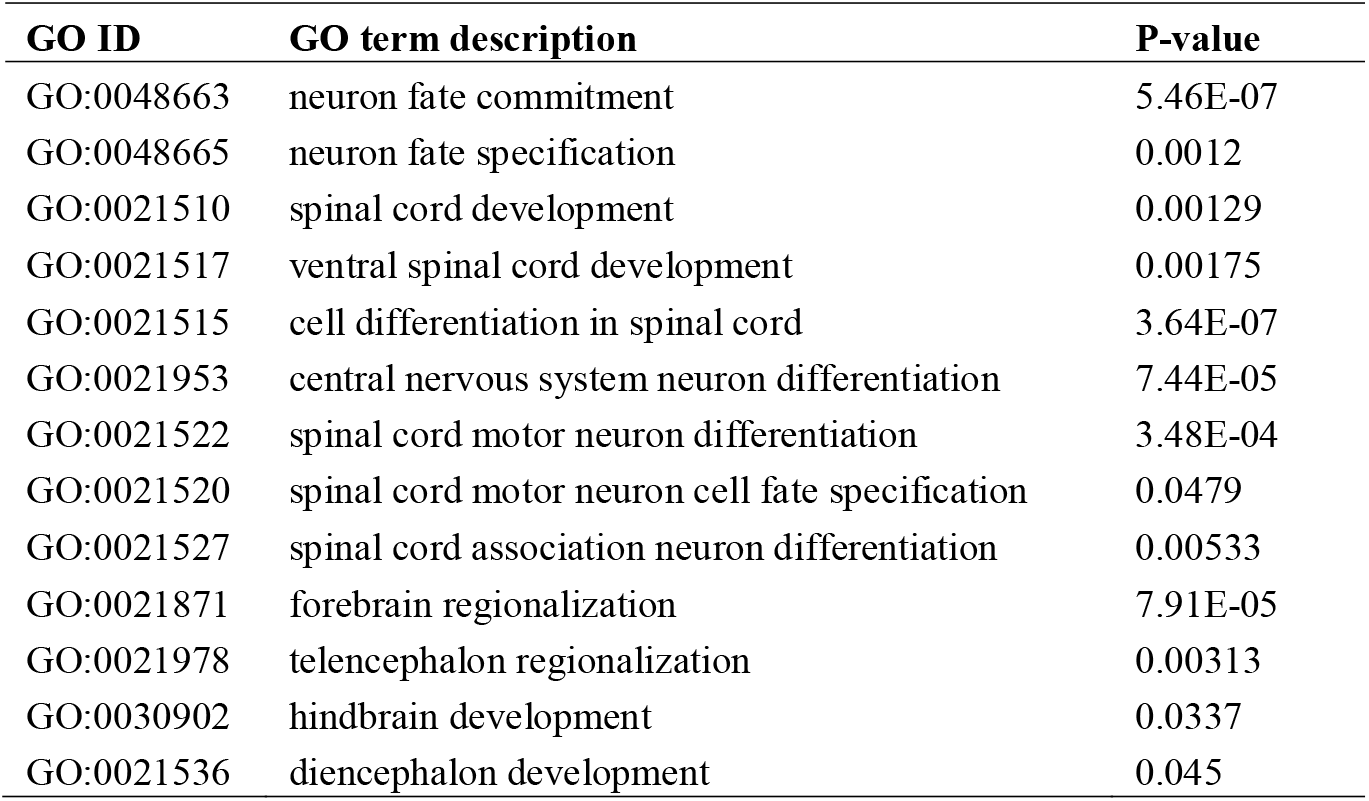
Significantly enriched nervous system-related GO terms for CNCRs at density of 0.3. P-value relates to the p-value for enrichment calculated using g:Profiler and its own g:SCS correction method^28^.

## WEB RESOURCES

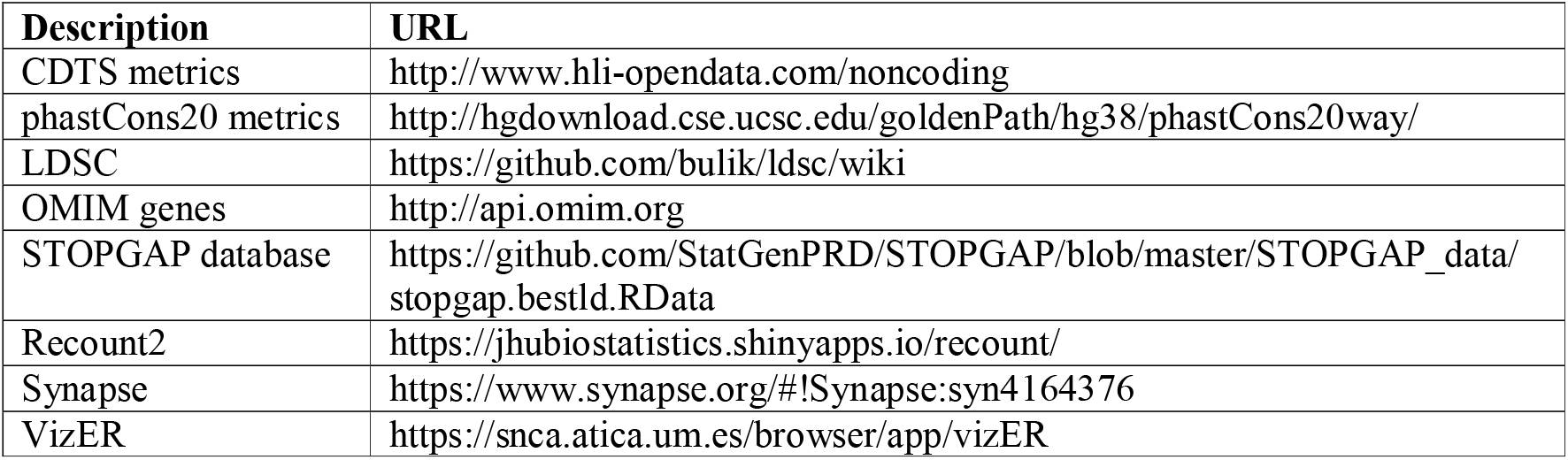

### International Parkinson’s Disease Genomics Consortium (IPDGC)

A. Noyce, A. Tucci, B. Middlehurst, D. Kia, M. Tan, H. Houlden, H. R. Morris, H. Plun-Favreau, P. Holmans, J. Hardy, D. Trabzuni, J. Bras, K. Mok, K. Kinghorn, N. Wood, P. Lewis, R. Guerreiro, R. Lovering, L. R’Bibo, M. Rizig, V. Escott-Price, V. Chelban, T. Foltynie, N. Williams, A. Brice, F. Danjou, S. Lesage, M. Martinez, A. Giri, C. Schulte, K. Brockmann, J. Simón-Sánchez, P. Heutink, P. Rizzu, M. Sharma, T. Gasser, A. Nicolas, M. Cookson, F. Faghri, D. Hernandez, J. Shulman, L. Robak, S. Lubbe, S. Finkbeiner, N. Mencacci, C. Lungu, S. Scholz, X. Reed, H. Leonard, G. Rouleau, L. Krohan, J. van Hilten, J. Marinus, A. Adarmes-Gómez, M. Aguilar, I. Alvarez, V. Alvarez, F. Javier Barrero, J. Bergareche Yarza, I. Bernal-Bernal, M. Blazquez, M. Bonilla-Toribio Bernal, M. Boungiorno, Dolores Buiza-Rueda, A. Cámara, M. Carcel, F. Carrillo, M. Carrión-Claro, D. Cerdan, J. Clarimón, Y. Compta, M. Diez-Fairen, O. Dols-Icardo, J. Duarte, R. l. Duran, F. Escamilla-Sevilla, M. Ezquerra, M. Fernández, R. Fernández-Santiago, C. Garcia, P. García-Ruiz, P. Gómez-Garre, M. Gomez Heredia, I. Gonzalez-Aramburu, A. Gorostidi Pagola, J. Hoenicka, J. Infante, S. Jesús, A. Jimenez-Escrig, J. Kulisevsky, M. Labrador-Espinosa, J. Lopez-Sendon, A. López de Munain Arregui, D. Macias, I. Martínez Torres, J. Marín, M. Jose Marti, J. Martínez-Castrillo, C. Méndez-del-Barrio, M. Menéndez González, A. Mínguez, P. Mir, E. Mondragon Rezola, E. Muñoz, J. Pagonabarraga, P. Pastor, F. Perez Errazquin, T. Periñán-Tocino, J. Ruiz-Martínez, C. Ruz, A. Sanchez Rodriguez, M. Sierra, E. Suarez-Sanmartin, C. Tabernero, J. Pablo Tartari, C. Tejera-Parrado, E. Tolosa, F. Valldeoriola, L. Vargas-González, L. Vela, F. Vives, A. Zimprich, L. Pihlstrom, P. Taba, K. Majamaa, A. Siitonen, N. Okubadejo & O. Ojo

## References

1 Walker, L. C. & Jucker, M. The Exceptional Vulnerability of Humans to Alzheimer’s Disease. Trends in molecular medicine 23, 534–545, doi:10.1016/j.molmed.2017.04.001 (2017).

2 O’Bleness, M., Searles, V. B., Varki, A., Gagneux, P. & Sikela, J. M. Evolution of genetic and genomic features unique to the human lineage. Nature reviews. Genetics 13, 853–866, doi:10.1038/nrg3336 (2012).

3 Xu, K., Schadt, E. E., Pollard, K. S., Roussos, P. & Dudley, J. T. Genomic and Network Patterns of Schizophrenia Genetic Variation in Human Evolutionary Accelerated Regions. Molecular Biology and Evolution 32, 1148–1160, doi:10.1093/molbev/msv031 (2015).

4 Cookson, M. R. Evolution of neurodegeneration. Current biology: CB 22, R753–761, doi:10.1016/j.cub.2012.07.008 (2012).

5 Diederich, N. J., James Surmeier, D., Uchihara, T., Grillner, S. & Goetz, C. G. Parkinson’s disease: Is it a consequence of human brain evolution? Movement disorders: official journal of the Movement Disorder Society, doi:10.1002/mds.27628 (2019).

6 Gearing, M., Rebeck, G. W., Hyman, B. T., Tigges, J. & Mirra, S. S. Neuropathology and apolipoprotein E profile of aged chimpanzees: implications for Alzheimer disease. Proceedings of the National Academy of Sciences of the United States of America 91, 9382-9386 (1994).

7 Collier, T. J., Kanaan, N. M. & Kordower, J. H. Ageing as a primary risk factor for Parkinson’s disease: evidence from studies of non-human primates. Nature reviews. Neuroscience 12, 359–366, doi:10.1038/nrn3039 (2011).

8 Raichlen, D. A. & Alexander, G. E. Exercise, APOE genotype, and the evolution of the human lifespan. Trends in neurosciences 37, 247–255, doi:10.1016/j.tins.2014.03.001 (2014).

9 Kellis, M. et al. Defining functional DNA elements in the human genome. Proceedings of the National Academy of Sciences of the United States of America 111, 6131–6138, doi:10.1073/pnas.1318948111 (2014).

10 Telenti, A. et al. Deep sequencing of 10,000 human genomes. Proceedings of the National Academy of Sciences of the United States of America 113, 11901–11906, doi:10.1073/pnas.1613365113 (2016).

11 di Iulio, J. et al. The human noncoding genome defined by genetic diversity. Nature genetics 50, 333–337, doi:10.1038/s41588-018-0062-7 (2018).

12 Lek, M. et al. Analysis of protein-coding genetic variation in 60,706 humans. Nature 536, 285–291, doi:10.1038/nature19057 (2016).

13 Schrider, D. R. & Kern, A. D. Inferring Selective Constraint from Population Genomic Data Suggests Recent Regulatory Turnover in the Human Brain. Genome biology and evolution 7, 3511–3528, doi:10.1093/gbe/evv228 (2015).

14 Ward, L. D. & Kellis, M. Evidence of abundant purifying selection in humans for recently acquired regulatory functions. Science (New York, N.Y.) 337, 1675–1678, doi:10.1126/science.1225057 (2012).

15 The Genomes Project, C. et al. A map of human genome variation from population-scale sequencing. Nature 467, 1061, doi:10.1038/nature09534 (2010).

16 Sherry, S. T. et al. dbSNP: the NCBI database of genetic variation. Nucleic Acids Res 29, 308–311 (2001).

17 Sudmant, P. H. et al. An integrated map of structural variation in 2,504 human genomes. Nature 526, 75, doi:10.1038/nature15394 (2015).

18 Siepel, A. et al. Evolutionarily conserved elements in vertebrate, insect, worm, and yeast genomes. Genome research 15, 1034–1050, doi:10.1101/gr.3715005 (2005).

19 Harrow, J. et al. GENCODE: the reference human genome annotation for The ENCODE Project. Genome research 22, 1760–1774, doi:10.1101/gr.135350.111 (2012).

20 Aken, B. L. et al. Ensembl 2017. Nucleic Acids Research 45, D635–D642, doi:10.1093/nar/gkw1104 (2017).

21 Finucane, H. K. et al. Partitioning heritability by functional annotation using genome-wide association summary statistics. Nature genetics 47, 1228–1235, doi:10.1038/ng.3404 (2015).

22 Bulik-Sullivan, B. K. et al. LD Score regression distinguishes confounding from polygenicity in genome-wide association studies. Nature genetics 47, 291–295, doi:10.1038/ng.3211 (2015).

23 Savage, J. E. et al. Genome-wide association meta-analysis in 269,867 individuals identifies new genetic and functional links to intelligence. Nature genetics 50, 912–919, doi:10.1038/s41588-018-0152-6 (2018).

24 Jansen, I. E. et al. Genome-wide meta-analysis identifies new loci and functional pathways influencing Alzheimer’s disease risk. Nature genetics 51, 404–413, doi:10.1038/s41588-018-0311-9 (2019).

25 Nalls, M. A. et al. Identification of novel risk loci, causal insights, and heritable risk for Parkinson’s disease: a meta-analysis of genome-wide association studies. The Lancet. Neurology 18, 1091–1102, doi:10.1016/s1474-4422(19)30320-5 (2019).

26 Pardinas, A. F. et al. Common schizophrenia alleles are enriched in mutation-intolerant genes and in regions under strong background selection. Nature genetics 50, 381–389, doi:10.1038/s41588-018-0059-2 (2018).

27 Wray, N. R. et al. Genome-wide association analyses identify 44 risk variants and refine the genetic architecture of major depression. Nature genetics 50, 668–681, doi:10.1038/s41588-018-0090-3 (2018).

28 Reimand, J. et al. g:Profiler-a web server for functional interpretation of gene lists (2016 update). Nucleic Acids Res 44, W83–89, doi:10.1093/nar/gkw199 (2016).

29 Supek, F., Bosnjak, M., Skunca, N. & Smuc, T. REVIGO summarizes and visualizes long lists of gene ontology terms. PloS one 6, e21800, doi:10.1371/journal.pone.0021800 (2011).

30 Hamosh, A., Scott, A. F., Amberger, J., Valle, D. & McKusick, V. A. Online Mendelian Inheritance in Man (OMIM). Human mutation 15, 57–61, doi: 10.1002/(sici) 1098-1004(200001)15:1<57::Aid-humu12>3.0.Co;2-g (2000).

31 Shen, J., Song, K., Slater, A. J., Ferrero, E. & Nelson, M. R. STOPGAP: a database for systematic target opportunity assessment by genetic association predictions. Bioinformatics 33, 2784–2786, doi:10.1093/bioinformatics/btx274 %J Bioinformatics (2017).

32 Human genomics. The Genotype-Tissue Expression (GTEx) pilot analysis: multitissue gene regulation in humans. Science (New York, N.Y.) 348, 648–660, doi:10.1126/science.1262110 (2015).

33 Bennett, D. A., Schneider, J. A., Arvanitakis, Z. & Wilson, R. S. Overview and findings from the religious orders study. Current Alzheimer research 9, 628–645, doi:10.2174/156720512801322573 (2012).

34 Collado-Torres, L. et al. Reproducible RNA-seq analysis using recount2. Nature Biotechnology 35, 319–321, doi:10.1038/nbt.3838 (2017).

35 Ng, B. et al. An xQTL map integrates the genetic architecture of the human brain’s transcriptome and epigenome. Nature neuroscience 20, 1418–1426, doi:10.1038/nn.4632 (2017).

36 Belloy, M. E., Napolioni, V. & Greicius, M. D. A Quarter Century of APOE and Alzheimer’s Disease: Progress to Date and the Path Forward. Neuron 101, 820–838, doi:10.1016/j.neuron.2019.01.056 (2019).

37 Mahley, Robert W. & Huang, Y. Apolipoprotein E Sets the Stage: Response to Injury Triggers Neuropathology. Neuron 76, 871–885 (2012).

38 Xu, Q. et al. Intron-3 retention/splicing controls neuronal expression of apolipoprotein E in the CNS. The Journal of neuroscience: the official journal of the Society for Neuroscience 28, 1452–1459, doi:10.1523/jneurosci.3253-07.2008 (2008).

39 Dieter, L. S. & Estus, S. Isoform of APOE with retained intron 3; quantitation and identification of an associated single nucleotide polymorphism. Mol Neurodegener 5, 34–34, doi:10.1186/1750-1326-5-34 (2010).

40 Zhang, D. et al. Incomplete annotation of disease-associated genes is limiting our understanding of Mendelian and complex neurogenetic disorders. 499103, doi:10.1101/499103 %J bioRxiv (2019).

41 Collado-Torres, L. et al. Flexible expressed region analysis for RNA-seq with derfinder. Nucleic Acids Res 45, e9, doi:10.1093/nar/gkw852 (2017).

42 Hindorff, L. A. et al. Potential etiologic and functional implications of genome-wide association loci for human diseases and traits. Proceedings of the National Academy of Sciences of the United States of America 106, 9362–9367, doi:10.1073/pnas.0903103106 (2009).

43 Sarropoulos, I., Marin, R., Cardoso-Moreira, M. & Kaessmann, H. Developmental dynamics of lncRNAs across mammalian organs and species. Nature 571, 510–514, doi:10.1038/s41586-019-1341-x (2019).

44 Cardoso-Moreira, M. et al. Gene expression across mammalian organ development. Nature 571, 505–509, doi:10.1038/s41586-019-1338-5 (2019).

45 Rathelot, J. A. & Strick, P. L. Subdivisions of primary motor cortex based on cortico-motoneuronal cells. Proceedings of the National Academy of Sciences of the United States of America 106, 918–923, doi:10.1073/pnas.0808362106 (2009).

46 de Noordhout, A. M. et al. Corticomotoneuronal synaptic connections in normal man: an electrophysiological study. Brain: a journal of neurology 122 (Pt 7), 1327–1340 (1999).

47 Al-Chalabi, A. et al. The genetics and neuropathology of amyotrophic lateral sclerosis. Acta neuropathologica 124, 339–352, doi:10.1007/s00401-012-1022-4 (2012).

48 Yu, J. T., Tan, L. & Hardy, J. Apolipoprotein E in Alzheimer’s disease: an update. Annual review of neuroscience 37, 79–100, doi:10.1146/annurev-neuro-071013-014300 (2014).

